# Linking Normative Models of Natural Tasks to Descriptive Models of Neural Response

**DOI:** 10.1101/158741

**Authors:** Priyank Jaini, Johannes Burge

## Abstract

Understanding how nervous systems exploit task relevant properties of sensory stimuli to perform natural tasks is fundamental to the study of perceptual systems. However, there are few formal methods for determining which stimulus properties are most useful for a given task. As a consequence, it is difficult to develop principled models for how to compute task-relevant latent variables from natural signals, and it is difficult to evaluate descriptive models fit to neural response. Accuracy Maxmization Analysis (AMA) is a recently developed Bayesian method for finding the optimal task-specific filters (receptive fields). Here, we introduce AMA-Gauss, a new faster form of AMA that incorporates the assumption that the class-conditional filter responses are Gaussian distributed. Next, we use AMA-Gauss to show that its assumptions are justified for two fundamental visual tasks: retinal speed estimation and binocular disparity estimation. Then, we show that AMA-Gauss has striking formal similarities to popular quadratic models of neural response: the energy model and the Generalized Quadratic Model (GQM). Together, these developments deepen our understanding of why the energy model of neural response have proven useful, improve our ability to evaluate results from subunit model fits to neural data, and should help accelerate psychophysics and neuroscience research with natural stimuli.

## 1. Introduction

Perceptual systems capture and process sensory stimuli to obtain information about behaviorally relevant properties of the environment. Characterizing the features of sensory stimuli and the processing rules that nervous systems use is central to the study of perceptual systems. Most sensory stimuli are high-dimensional, but only a small set of stimulus features are relevant for any particular task. Thus, perceptual and neural processing in particular tasks is driven by sets of features that occupy a lower dimensional space (i.e. can be described more compactly) than the stimuli themselves. These considerations have motivated perception and neuroscience researchers to develop methods for dimensionality reduction that characterize the statistical properties of proximal stimuli, that describe the responses of neurons to those stimuli, and that specify how those responses could be decoded [3, 11, 12, 24, 28, 37, 43, 49, 45, 44, 50, 40, 31, 39]. However, many of these methods are task-independent; that is, they do not explicitly consider the sensory, perceptual, or behavioral tasks for which the encoded information will be used. Empirical studies in psychophysics and neuroscience often focus on the behavioral limits and neurophysiological underpinnings of performance in specific tasks. Thus, there is a partial disconnect between task-independent theories of encoding and common methodological practices in psychophysics, and sensory and systems neuroscience.

Task-specific normative models prescribe how best to perform a particular task. Task-specific normative models are useful because they provide principled hypotheses about i) the stimulus features that nervous systems should encode and ii) the processing rules that nervous systems should use to decode the encoded information. Normative models in widespread use are often not directed at specific tasks. Methods for fitting neural response cannot generally be interpreted with respect to specific tasks. Accuracy Maximization Analysis (AMA) is a Bayesian method for finding the stimulus features that are most useful for specific tasks [21, 8]. In conjunction with carefully calibrated natural stimulus databases, AMA has contributed to the development of normative models of several fundamental tasks in early- and mid-level vision [4, 5, 6, 7], by determining the encoding filters (receptive fields) that support optimal performance in each task. These normative models have, in turn, predicted major aspects of primate neurophysiology and human psychophysical performance with natural and artificial stimuli [6, 7].

The primary theoretical contribution of this manuscript is to establish formal links between normative models of specific tasks and popular descriptive models of neural response (Fig. 1). To do so, we first develop a new form of AMA called AMA-Gauss, which incorporates the assumption that the latent-variable-conditioned filter responses are Gaussian distributed. Then, we use AMA-Gauss to find the filters (receptive fields) and pooling rules that are optimal with natural stimuli for two fundamental tasks: estimating the speed of retinal image motion and estimating binocular disparity [6, 7]. For these two tasks, we find that the critical assumption of AMA-Gauss is justified: the optimal filter responses to natural stimuli, conditioned on the latent variable (i.e. speed or disparity), are indeed Gaussian distributed. Then, we show that this empirical finding provides a normative explanation for why neurons that select for motion and disparity have been productively modeled with energy-model-like (i.e. quadratic) computations [34, 35, 36, 13, 14]. Finally, we recognize and make explicit the formal similarities between AMA-Gauss and the Generalized Quadratic Model (GQM) [40, 54], a recently developed method for neural systems identification. These advances may help bridge the gap between empirical studies of psychophysical and neurophysiological tasks, methods for neural systems identification, and task-specific normative modeling (Fig. 1).

**Figure 1:**
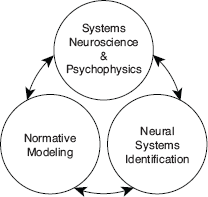
Linking scientific subfields. Perception science benefits when links are drawn between psychophysical and neuroscience studies of particular tasks, task-agnostic statistical procedures that fit models to data, and task-specific normative methods that determine which models are best. The current work develops formal links between the energy model for describing neural response, the Generalized Quadratic Model (GQM) for fitting neural response, and AMA-Gauss for determining the neural response properties that best serve a particular task.

In addition to these theoretical contributions, the development of AMA-Gauss represents a technical advance. The major drawback of AMA is its computational expense. Its compute-time for filter learning is quadratic in the number of stimuli in the training set, rendering the method impractical for large-scale problems without specialized computing resources. We demonstrate, both analytically and empirically, that AMA-Gauss reduces compute-time for filter learning from quadratic to linear. Thus, for tasks for which the critical assumption of AMA-Gauss is justified, AMA-Gauss can be of great practical benefit.

### Background

#### Energy Model

Energy models have been remarkably influential in visual neuroscience. The standard energy model posits two Gabor-shaped subunit receptive fields, the responses of which are squared and then summed (Fig. 2.A). These computations yield decreased sensitivity to the local position of stimulus features (i.e. spatial phase) and increased sensitivity to task-relevant latent variable. Energy models have been widely used to describe the computations of neurons involved in coding retinal image motion and binocular disparity [1, 13, 14]. However, the motion-energy and disparity-energy computations are primarily descriptive models of a neuron’s response properties. The energy model does not make explicit how neural responses should be decoded into estimates.

**Figure 2:**
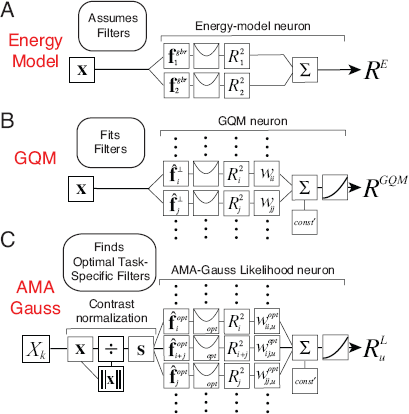
Computations of an energy model neuron, a GQM model neuron, and an AMA-Gauss likelihood neuron. All three have quadratic computations at their core. The energy model and the GQM describe the computations that neurons perform. AMA-Gauss prescribes the computations that neurons should perform to optimize performance in a specific task. **A** The standard energy model assumes two Gabor-shaped orthogonal subunit filters (receptive fields) **f**^*gbr*^ to account for a neuron’s response. The response of an energy model neuron *R^E^* is obtained by adding the squared responses of the filters. **B** The GQM fits multiple arbitrarily-shaped orthogonal subunit receptive fields **f**^⊥^ that best account for a neuron’s response. The response of a GQM model neuron *R^GQM^* is obtained by pooling the squared (and linear, not shown) responses of the subunit filters via a weighted sum, and passing the sum through an output nonlinearity. **C** AMA-Gauss finds the optimal subunit filters **f**^*opt*^ and quadratic pooling rules for a specific task. Unlike standard forms of the energy model and the GQM, AMA-Gauss incorporates contrast normalization and finds subunit filters that are not necessarily orthogonal. The response 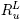 of an AMA-Gauss likelihood neuron represents the likelihood of latent variable *X_u_*. The likelihood is obtained by pooling the squared (and linear, not shown) subunit filter responses, indexed by *i* and *j*, via a weighted sum (Eq. (20)).

Under what circumstances would energy-model-like computations be optimal? Energy-model-like computations would be optimal if quadratic pooling is necessary for determining the likelihood of the task-relevant latent variable. We show below that for retinal speed and binocular disparity estimation, two tasks classically associated with the energy model, quadratic pooling is indeed necessary to optimally decode the task-relevant latent variable (see Section 3). Therefore energy-model-like computations are optimal for these tasks. AMA-Gauss is specifically designed to find the receptive fields and pooling rules that optimize performance under these conditions. It is thus likely to help accelerate the development of normative models of other tasks for which the energy model has provided a useful description.

#### Generalized Quadratic Model (GQM)

The standard energy model assumes that the responses of certain neurons can be accounted for by two Gabor-shaped subunit receptive fields. Real neurons are not constrained to have only two subunit receptive fields, nor are their shapes constrained to be Gabor-shaped. The Generalized Quadratic Model (GQM) fits multiple arbitrarily-shaped subunit filters and quadratic pooling rules that best account for a neuron’s response (Fig. 2.B; [40]). The GQM is a specific example of a large class of models designed for neural systems identification, collectively known as ’subunit models’. The spike triggered average (STA), spike triggered covariance (STC), and the generalized linear model (GLM) are popular examples of this class of models. The goal of these models is to provide a computational level description of a neuron’s computations that can predict its responses to arbitrary stimuli.

Unfortunately, a tight description of a neuron’s computations does not necessarily provide insight about how (or whether) that neuron and its computations subserve a specific task; after a subunit model has been fit, the purpose of the neuron’s computations is often unclear. Thus, although methods for neural systems identification are essential for determining what the components of nervous systems do, they are unlikely to determine why they do what they do. One way to address this issue is to develop normative frameworks (i) that determine the computations that are optimal for particular tasks and (ii) that share the same or similar functional forms as popular methods for describing neural response.

AMA-Gauss is a normative method that is designed to find the filters (receptive fields) and quadratic pooling rules that are optimal for specific sensory-perceptual tasks (Fig. 2.C; see 2). AMA-Gauss has a functional form that is closely related to the energy model and the GQM, but it has a different aim. Rather than describing what a neuron does, it prescribes what neurons should do. In fact, given a hypothesis about the function of a particular neuron, AMA-Gauss can predict the subunit filters and pooling rules that will be recovered by the GQM. The development of closely related normative models and methods for neural systems identification is likely to enhance our ability to interpret fits to neural data and accelerate progress in psychophysical and neuroscientific research.

## 2. Methods

This section formally develops AMA-Gauss. To provide context for this technical contribution, we first review the setup and main equations for AMA [21]. Then, we derive the main equations for AMA-Gauss, provide a geometric intuition for how it works, and discuss practices for best use. Readers that are more interested in the scientific implications, and less interested in the mathematical formalisms, can skip ahead to Results.

### Accuracy Maximization Analysis

The goal of AMA is to find the filters (receptive fields) that extract the most useful stimulus features for a particular task. Consistent with real biological systems, AMA filters are corrupted by response noise, and it places no constraints on the orthogonality of its filters. AMA searches for the optimal filters with a closed form expression for the cost that relies on a Bayes optimal decoder. The filters are constrained to have unit magnitude 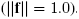. The expression for the cost requires the specification of five factors (see Fig 3.A). These factors are i) a well defined task (i.e. a latent variable to estimate from high-dimensional stimuli), ii) a labeled training set of stimuli, iii) a set of encoding filters, iv) a response noise model, v) and a cost function (Fig. 3.A). The training set specifies the joint probability distribution 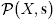 between the latent variable *X* and the stimuli **s** (Fig. 3.B) and implicitly defines the prior 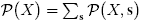 over the latent variable (see Discussion). If the training set is representative, results will generalize well to stimuli outside the training set.

**Figure 3:**
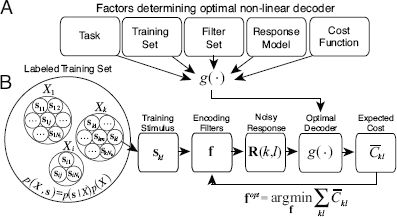
Accuracy Maximization Analysis. **A** Factors determining the optimal decoder used by AMA during filter learning. **B** Steps for finding optimal task-specific filters via AMA.

For any particular filter set, the matched Bayes optimal decoder provides the cost by computing the posterior probability over the latent variable 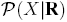, reading out the optimal estimate from the posterior, and then assigning a cost to the error. The steps for finding the optimal task-specific filters are: i) select a particular stimulus **s**_*kl*_ from the labeled training set, ii) obtain a set of noisy filter responses **R** (*k*, *l*) from a particular (possibly non-optimal) set of filters, iii) use the optimal non-linear decoder g(.) to obtain the optimal estimate 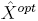 and its expected cost 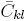, iv) repeat for each stimulus and compute the average cost across all stimuli in the training set, v) update the filters to reduce the cost, vi) repeat until the average cost is minimized. The optimal task-specific filters **f**^*opt*^ are those that minimize the cost (Fig. 3.B).

### Bayes Optimal Decoder and Filter Response Model

The Bayes optimal decoder gives a closed form expression for the cost for any filter or set of filters, given the training stimuli. The posterior probability of latent variable *X_u_* given the noisy filter responses **R** (*k*, *l*) to stimulus **s**_*kl*_ is given by Bayes’ rule

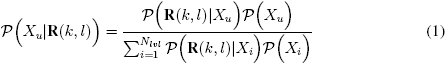

where *N_lvl_* is the number of latent variable level, and *l* indexes the stimuli having latent variable value *X_k_*. The conditional distribution of noisy responses given the latent variable is

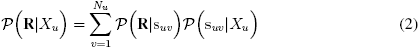

where *N_u_* is the number of stimuli having latent variable level *X_u_*, and *v* indexes training stimuli having that latent variable value. Conveniently, 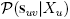 and 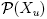 are determined by the training set; 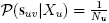 is the probability of particular stimulus *v* with Nulatent variable *X_u_* given that there are *N_u_* such stimuli, and 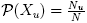 is the fraction of all stimuli having latent variable *X_u_*. Therefore, Equation 1 reduces to

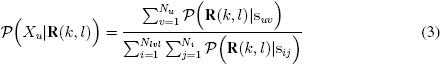

Equation 3 indicates that the posterior probability is given by the sum of the within-level stimulus likelihoods, normalized by the sum of all stimulus likelihoods.

Our aim is to understand task-specific information processing in biological systems. Thus, the response noise model should be consistent with the properties of biological encoders. AMA uses scaled additive (e.g. Poisson-like) Gaussian noise, a broadly used model of neural noise in early visual cortex [20]. Equations 4-7 define the response model, and specify the distribution of noisy filter responses 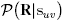 to each stimulus. For an individual filter **f**_*t*_ from set of filters **f** =[**f**_1_,**f**_2_,…,**f**_*q*_] (where *q* is the number of filters), the mean response *r_t_*, noisy response *R_t_*, and noise variance 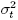 to stimulus **s**_*uv*_ having latent variable value *X_u_* are

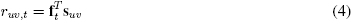

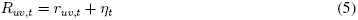

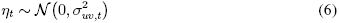

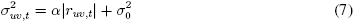

where η is a noise sample, α is the fano factor, and 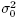 is baseline noise variance. The proximal stimulus 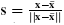 is contrast-normalized consistent with standard models [2, 23], where **x** is a (possibly noise corrupted) intensity stimulus. If *q* filters are considered simultaneously, the response distributions 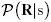, and the variables in Equations 4-7 become *q*-dimensional: mean response vector **r** = [*r*_1_, *r*_2_, …, *r_q_*], noisy response vector **R** = [*R*_1_, *R*_2_, …, *R_q_*], and response noise covariance matrix Λ.

The posterior probability distribution over the latent variable given the noisy filter responses to any stimulus in the training set is fully specified by Equations 3-7. The next step is to define the cost associated with a noisy response to an individual stimulus. The cost is given by

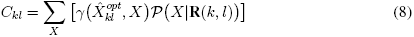

where γ(.) is an arbitrary cost function and 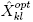 is the optimal estimate associated with noisy response **R** (*k*, *l*). The overall cost for a set of filters is the expected cost for each stimulus averaged over all stimuli

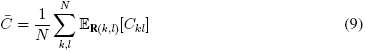

The goal of AMA is to obtain the filters **f** that minimize the overall cost

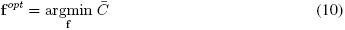

where **f**^*opt*^ are the optimal filters.

A single evaluation of the posterior probability distribution (Eq. 3) for each stimulus in the training set requires 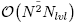 operations where *N* is the total number of stimuli and *N_lvl_* is the number of latent variable levels in the training set. As noted earlier, this compute-time makes AMA impractical for large scale problems without specialized computing resources.

There are at least two methods for achieving significant computational savings in optimization problems: employing models with strong parametric assumptions, and employing stochastic gradient descent routines. Both methods have drawbacks. Models with strong parametric assumptions are only appropriate for cases in which the assumptions approximately hold. Stochastic gradient descent routines are noisy and may not converge to the optimum filters. We have previously developed AMA-SGD, a stochastic gradient descent routine for AMA [8]. Here, we develop AMA-Gauss, a model with strong parametric assumptions.

### AMA-Gauss

In this section, we first introduce AMA-Gauss and highlight its advantages over AMA. Subsequently, we provide expressions for AMA-Gauss likelihood function, *L*_2_ and *L*_0_ cost functions, and their gradients. We believe this is a valuable step towards making AMA a more practical tool in vision research.

#### AMA-Gauss: Class-conditional Gaussian Assumption

AMA-Gauss is a version of AMA that makes the parametric assumption that the filter responses are Gaussian distributed when they are conditioned on a particular value of the latent variable

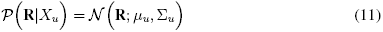

where **R** are responses to stimuli having latent varible level *X_u_*,

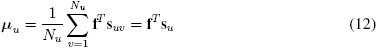

is the class-conditional mean of the noisy filter responses and

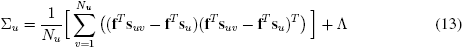

is the class-conditional covariance of the noisy filter responses. The first term in Eq. 13 is the class-conditional covariance of the expected filter responses. The second term in Eq. 13, Λ, is the covariance matrix of the filter response noise 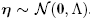. There are two major reasons for making the Gaussian assumption. First, if the response distributions are Gaussian, then AMA-Gauss will return the same filters as AMA while simultaneously providing huge savings in compute-time. Second, the assumption is justified for at least two fundamental visual tasks in early vision (see Results; [6, 7]). With time, we speculate that similar statements will be justified for other sensory-perceptual tasks.

Under the AMA-Gauss assumption, the posterior probability (Eq 1) of latent variable *X_u_* is

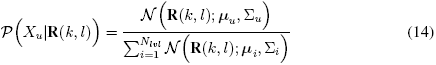

where *N_lvl_* is the number of latent variable levels. The AMA-Gauss posterior (Eq 14), has a simpler form than the AMA posterior (Eq 3). Hence, whereas a single evaluation of the AMA posterior probability distribution requires 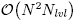 perations (Eq 3), the AMA-Gauss posterior requires only 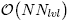 operations where *N* is the number of stimuli in the training set (see Section 3). This reduction in compute-time substantially improves the practicality of AMA when the Gaussian assumption is justified. Even if the Gaussian assumption is not justified, AMA-Gauss is guaranteed to make the best possible use of first- and second-order conditional response statistics, and could thus provide a decent initialization at low computational cost.

#### AMA-Gauss: Derivations of the Likelihood Function, Costs, and Gradients

Analytic solutions for the optimal filters under AMA-Gauss (and AMA) are not available in closed form. Here, we provide expressions for the AMA-Gauss likelihood function, *L*_2_ cost, *L*_0_ cost, and their gradients.

The maximum likelihood AMA-Gauss encoding filters 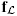 are those that simultaneously maximize the likelihood of the correct latent variable *X_k_* across all stimuli in the training set. Stimuli having latent variable value *X_k_* are indexed by *l*, and the *i^th^* stimulus in the training set is denoted (*k_i_*, *l_i_*). The likelihood function of the AMA-Gauss filters is

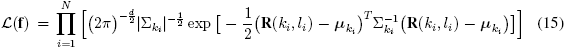

The maximum likelihood filters can be determined by maximizing the likelihood function or, equivalently, minimizing the negative log-likelihood function

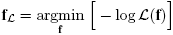

In practice, the expected negative log-likelihood is easier to minimize. Complete derivations of the likelihood function, the expected log-likelihood function, and closed form expressions for the associated gradients are provided in Appendix A. These expressions can be used to estimate the maximum-likelihood filters via gradient descent.

Next, we derive the AMA-Gauss cost for two popular cost functions for which the minimum mean squared error (MMSE) estimate and maximum a posteriori (MAP) are optimal: the *L*_2_ and *L*_0_ cost. The cost function specifies the penalty assigned to different types of error. For the *L*_2_ (i.e. squared error) cost function, the expected cost for each stimulus **s**_*kl*_(Eq 9) is

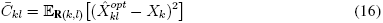

where the optimal estimate 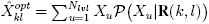 is the mean of the posterior.

For the *L*_0_ (i.e. 0,1) cost function, the expected cost across all stimuli is closely related to the KL-divergence of the observed posterior and an idealized posterior with all its mass at the correct latent variable *X_k_*; in both cases, cost is determined only by the posterior probability mass at the correct level of the latent variable [21, 8]. Here, the expected KL-divergence per stimulus is equal to the negative log-posterior probability at the correct level [21]

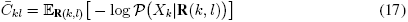

In a slight abuse of terminology, we refer to this divergence as the *L*_0_ or KL-divergence cost.

The gradient of the total expected cost across all stimuli can be evaluated by calculating the gradient of the cost for each stimulus 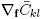 (see Eq (9)). Hence, the gradient of the total expected cost is

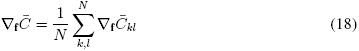

The gradient of the cost for each stimulus can be evaluated by calculating the gradient of the posterior probability. Complete derivations of the cost and the gradient of the cost for the *L*_2_ and *L*_0_ cost functions are given in Appendix B and Appendix C respectively.

Cost is minimized when responses to stimuli having different latent variable values overlap as little as possible. The cost functions (i.e. max-likelihood, *L*_0_ cost, *L*_2_ cost) exert pressure on the filters to produce class-conditional response distributions that are as different as possible given the constraints imposed by the stimuli. Hence the optimal filters will i) maximize the differences between the class-conditional means or covariances and ii) minimize the generalized variance for each class-conditional response distribution. (Generalized variance is a measure of overall scatter, represents the squared volume of the ellipse, and is given by the determinant of the covariance matrix.)

#### AMA-Gauss: Geometric Intuition

Fig. 4 provides a geometric intuition for the relationship between the filter response distributions, the likelihood, and the posterior probability distribution for two simple hypothetical cases. Both cases have three latent variable values. In one case, the information about the latent variable is carried by the class-conditional mean (Fig. 4.A-C). In the other case, the information about the latent variable is carried by the class-conditional covariance (Fig. 4.D-F). In all cases, the class-conditional responses to stimuli having the same latent variable value are Gaussian distributed. With a single filter, the response distributions are one-dimensional (Fig. 4.A,D). For any observed noisy response *R*, the likelihood of a particular level of the latent variable *X_u_* is found by evaluating its response distribution at the observed response (blue dot; Fig. 4.A,D). The posterior probability of latent variable *X_u_* is obtained by normalizing with the sum of the likelihoods (blue, red, and green dots; Fig. 4.B,E). With two filters, the response distributions are two-dimensional (red, blue, and green ellipses with corresponding marginals; Fig. 4.C,F). The second filter increases the posterior probability mass at the correct value of the latent variable (not shown) because the second filter selects for useful stimulus features that the first filter does not. These hypothetical cases illustrate why cost is minimized when mean or covariance differences are maximized between classes and generalized variance is minimized within classes. The filters that make the response distributions as different as possible make it as easy as possible to decode the latent variable.

**Figure 4:**
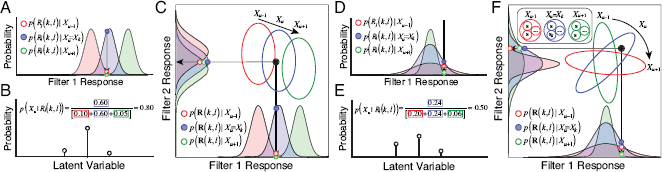
Relationship between conditional filter response distributions, likelihood, and posterior probability. Two hypothetical cases are considered, each with three latent variable values. **A** One-dimensional (i.e. single filter) Gaussian conditional response distributions, when information about the latent variable is carried only by the class-conditional mean; distribution means, but not variances, change with the latent variable. The blue distribution represents the response distribution to stimuli having latent variable value *X_k_*. The red and green distributions represent response distributions to stimuli having different latent variables values *X_u_* ≠ *X_k_*. The blue dot represents the likelihood 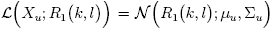 that observed noisy filter response *R*_1_ (*k*, *l*) to stimulus **s**_*k,l*_ was elicited by a stimulus having latent variable level *X_u_* = *X_k_*. Red and green dots represent the likelihoods that the response was elicited by a stimulus having latent variable *X_u_* ≠ *X_k_* (i.e. by a stimulus having the incorrect latent variable value). **B** Posterior probability over the latent variable given the noisy observed response in A. The posterior probability of the correct latent variable value (in this case, *X_k_*) is given by the likelihood of the correct latent variable value normalized by the sum of all likelihoods. Colored boxes surrounding entries in the inset equation indicate the likelihood of each latent variable. **C** Two-dimensional (i.e. two filter) Gaussian response distributions. Each ellipse represents the joint filter responses to all stimuli having the same latent variable value. The second filter improves decoding performance by selecting for useful stimulus features that the first filter does not. The black dot near the center of the blue ellipse represents an observed noisy joint response **R** (*k*, *l*)to stimulus **s**_*k,l*_. The likelihood 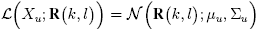 that the observed response was elicited by a stimulus having latent variable value *X_u_* is obtained by evaluating the joint Gaussian at the noisy response; in this case, the product of the likelihoods represented by the blue dots on the single filter response distributions. **D-F** Same as A-C, but where information about the latent variable is carried by the class-conditional covariance instead of the mean; ellipse orientation, but not location, changes with the latent variable. AMA-Gauss finds the filters yielding conditional response distributions that are as different from each other as possible, given stimulus constraints.

#### AMA-Gauss: Best Practices

The AMA-Gauss method developed here does not automatically determine the number of stimuli to train on, or the number of task-specific filters to learn; these choices are left to the user.

To obtain representative results (i.e. to minimize sampling error) the training set must be of sufficient size. AMA-Gauss uses the sample mean and covariance to approximate the Gaussian distributions of filter responses conditional on each value of the latent variable (Eq. (11)). Training sets with at least 250 stimuli per level tend to give representative results.

To extract all task-relevant information from each stimulus a sufficient number of receptive fields must be learned. In general, the best practice is to learn filters until the change in the value of the total cost is negligible [21]. The current paper aims to demonstrate the properties and usefulness of AMA-Gauss rather than determine the best number of filters; for clarity, we show only four filters for each task (see Section 3). Previous work has shown that, for the two tasks considered here, eight filters are required to capture nearly all task-relevant information [6, 7]. The results presented in this paper hold for all eight filters, but we show only four for ease of presentation.

## 3. Results

Retinal speed estimation and binocular disparity estimation are canonical visual tasks. Accurate and precise estimation of retinal image motion is critical for the accurate estimation of object motion and self motion through the environment. Accurate and precise estimation of binocular disparity is critical for the accurate estimation of depth and the control of fixational eye movements. Although of fundamental importance for mobile seeing organisms, both tasks are difficult in natural conditions because of the enormous variability and complexity in natural images.

The plan for the results section is as follows. First, we use AMA-Gauss^1^ to find the receptive fields that are optimal for estimating speed and disparity from local patches of natural images. Second, we compare AMA-Gauss and AMA and show that both methods (i) learn the same filters and (ii) converge to the same cost for both tasks. Third, we verify that AMA-Gauss achieves the expected reductions in compute-time: filter-learning with AMA-Gauss is linear whereas AMA is quadratic in the number of stimuli in the training set. Fourth, we show that the class-conditional filter responses are approximately Gaussian, thereby justifying the Gaussian assumption for these tasks. Fifth, we show how contrast normalization contributes to the Gaussianity of the class-conditional responses. Sixth, we explain how the filter response distributions determine the likelihood functions and optimal pooling rules. Seventh, we explain how these results provide a normative explanation for why energy-model-like computations describe the response properties of neurons involved in these tasks. Eighth, and last, we establish the formal relationship between AMA-Gauss and the GQM, a recently developed method for neural systems identification.

For each task, we obtained an existing labeled training set of natural photographic stimuli consisting of approximately ten thousand randomly sampled stimuli. All stimuli subtended 1 deg of visual angle. Perspective projection, physiological optics, and the wavelength sensitivity, spatial sampling, and temporal integration functions of the foveal cones were accurately modeled. Input noise was added to each stimulus with a noise level just high enough to mask retinal image detail that would be undetectable by the human visual system [53]. Both training sets had flat prior probability distributions 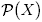 over the latent variable (see Discussion). The training set for speed estimation consisted of 10500 stimuli (10500 stimuli=500 stimuli/level x 21 levels; [7]). Retinal speeds ranged from −8 deg/sec to +8 deg/sec; negative and positive speeds correspond to leftward and rightward drifting movies. Each stimulus had a duration of 250ms. The training set for disparity estimation consisted of 7600 stimuli (7600 stimuli=400 stimuli/level x 19 levels; [6]). Binocular disparities ranged from −16.875 arcmin to +16.875 arcmin; negative and positive disparities correspond to uncrossed and crossed disparities. (Note that although these training sets have a discrete number of latent variable values, AMA filters can be learned with discrete- or with real-valued latent variables.) For extensive additional details on these training sets and for ideal observer performance in these tasks, please see [6, 7]. One important limitation of these datasets is that all motion signals were rigid and that all disparity signals were planar. Future work will examine the impact of non-rigid motion (e.g. looming) and local depth variation (e.g. occlusion) on performance (see Discussion).

Before processing, retinal image stimuli for both tasks were vertically averaged under a raised cosine window (0.5° at half-height). Vertically oriented receptive fields respond identically to the original and vertically averaged stimuli, and canonical receptive fields for both tasks are vertically oriented [6, 7]. Thus, the vertically averaged signals represent the signals available to the the orientation column that would be most useful to the task. Future work will examine the impact of off-vertical image features on performance.

Next, we used AMA-Gauss to find the optimal filters for both tasks. The results presented below were obtained using the *L*_0_ cost function and constant, additive, independent filter response noise. In general, we have found that the optimal filters are quite robust to the choice of cost function when trained with natural stimuli [8]. Figure 5 shows results for the retinal speed estimation task and Figure 6 shows results for the disparity estimation task. AMA-Gauss and AMA learn nearly identical encoding filters (Fig. 5.A and 6.A; ρ > 0.96) and exhibit nearly identical estimation costs (Fig. 5.B and 6.B). AMA-Gauss also dramatically reduces compute time (Fig. 5.CD and 6.CD). With AMA, the time required to learn filters increases quadratically with their number of stimuli in the training set. With AMA-Gauss, filter learning time increases linearly with the number of stimuli. Finally, the class-conditional filter responses are approximately Gaussian (Fig. 5EF and 6EF), indicating that the Gaussian assumption is justified for both tasks. Thus, quadratic computations are required to determine the likelihood of a particular value of the latent variable (see below). After characterizing the conditional response distributions, the posterior probability distribution over the latent variable 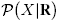 can be obtained by straightforward application of Bayes’ rule.

**Figure 5:**
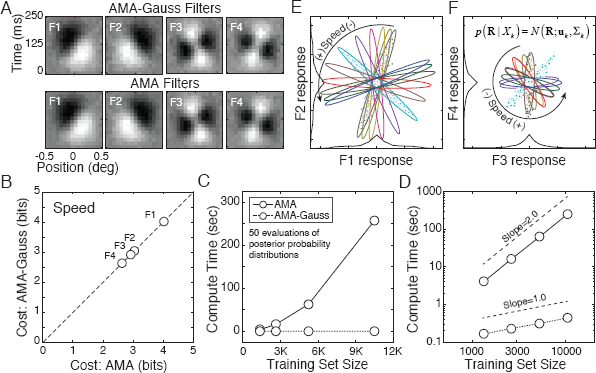
Speed estimation task: filters, cost, compute time, and class-conditional response distributions. **A** AMA-Gauss and AMA filters for estimating speed (−8 to +8 deg/sec) from natural image movies are nearly identical; ρ > 0.96 for all filters. **B** The cost for all the filters for both the models is identical. **C** Compute time for fifty evaluations of the posterior probability distribution is linear with AMA-Gauss, and quadratic with the full AMA model, in the training set size. **D** Same data as in C but on log-log axes. **E,F** Joint filter responses, conditioned on each level of the latent variable, are approximately Gaussian (also see below). Different colors indicate different speeds. Individual symbols represent responses to individual stimuli. Thin black curves show that the filter response distributions, marginalized over the latent variable 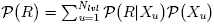, are heavier-tailed than Gaussians (see Sections 3, 4)

**Figure 6:**
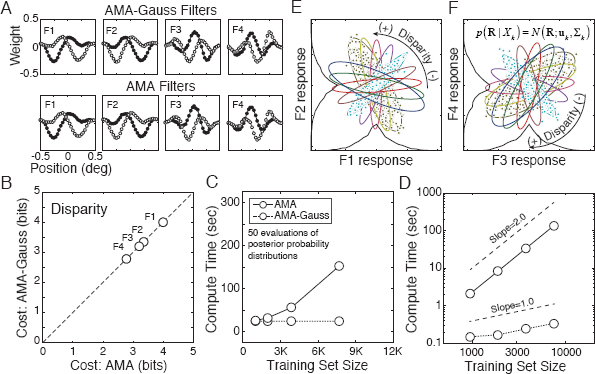
Disparity estimation task: filters, cost, compute time, and class-conditional response distributions. **A** AMA-Gauss and AMA filters for estimating disparity from natural stereo-images (−15 to +15 arcmin). **B-F** Caption format same as Fig. 5.B-F.

### Response Normalization, Response Gaussianity, and Decoding Performance

Contrast varies significantly in natural stimuli. How does contrast normalization affect the filter responses? For the class of problems considered here (e.g. retinal speed estimation, binocular disparity estimation, and other energy-model-related tasks), neurophysiologically plausible contrast normalization [2, 23] must be built into the filter response model (Eq. 4) for the class-conditional filter responses 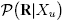 to be Gaussian distributed. (Note that many different models of normalization are computationally equivalent [2, 23].) In AMA-Gauss, the input stimulus **s** is a contrast normalized 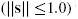 version of a (possibly noise-corrupted) intensity stimulus **x** with mean intensity 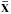. Luminance normalization converts the intensity stimulus to a contrast stimulus 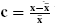 by subtracting off and dividing by the mean. Contrast normalization converts the contrast stimulus to a contrast normalized signal with unit magnitude (or less) 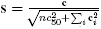 where *c*_50_ is an additive constant and *n* is the dimensionality of (e.g. number of pixels defining) each stimulus. Here, we assumed that the value of the additive constant is *c*_50_ = 0.0. The effect of the value of *c*_50_ has been studied previously [6].

To examine the effect of contrast normalization on the class-conditional filter response distributions, we computed the filter responses to the same stimuli with and without contrast normalization. With contrast normalization, filter response distributions are approximately Gaussian (Fig. 7.A,B,E,F). Without contrast normalization, filter response distributions have tails much heavier than Gaussian (Fig. 7.C,D,G,H). (Note that AMA-Gauss learns very similar filters with and without contrast normalization. Normalization does not change which stimulus features should be selected; it changes only how the selected features are represented.) Thus, biologically realistic normalization helps Gaussianize the conditional response distributions. Related results have been reported by other groups [51, 29].

**Figure 7:**
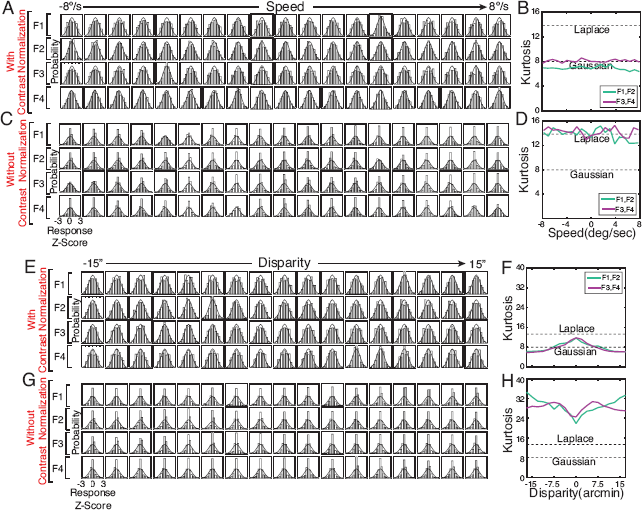
Filter responses with and without contrast normalization. **A** Class-conditional filter response distributions 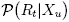 to contrast-normalized stimuli for each individual filter and each level of the latent variable in the speed estimation task. For visualization, responses are transformed to *Z*-scores by subtracting off the mean and dividing by the standard deviation. Gaussian probability density is overlaid for purposes of comparison. **B** Kurtosis of the two-dimensional conditional response distributions from filters 1 and 2 (violet; also see Fig. 5.E) and filters 3 and 4 (green; also see Fig. 5.F) for all levels of the latent variable. A two-dimensional Gaussian has a kurtosis of 8.0. Kurtosis was estimated by fitting a multidimensional generalized Gaussians via maximum likelihood methods. **C,D** Same as A,B but without contrast normalization. **E-H** Same as A-D, but for the task of disparity estimation.

Contrast normalization not only Gaussianizes the response distributions; it also improves performance. If response distributions are heavy-tailed and have strong peaks at zero, then the Gaussian assumption is violated and attempts to decode the latent variable from those responses suffer. Contrast normalization reduces the peak at zero, thereby reducing decoding difficulty. Fig.8 compares decoding cost in the speed and disparity tasks with and without contrast normalization and shows that failing to normalize harms performance. Thus, contrast normalization improves task performance by decreasing kurtosis and increasing response Gaussianity.

**Figure 8:**
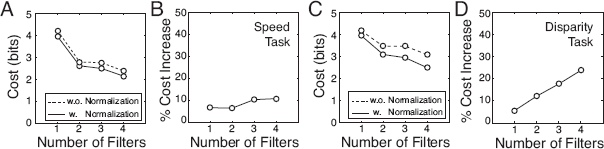
Decoding performance with and without contrast normalization. **A** Contrast normalization decreases decoding cost for the speed estimation task. **B** Percentage increase in cost without contrast normalization. **C-D** Same as A-B, but for the disparity estimation task. The same result holds for different cost functions (e.g. squared error) and larger number of filters. If eight filters are used in the disparity task, failing to contrast normalize can decrease performance by ~ 40%.

Subunit response models (e.g. the standard energy model, the GQM, and other LN models) are widely used to describe and fit neurons. They do not generally incorporate normalization (see below; [1, 44, 40, 50]). This fact is unsurprising. Many laboratory experiments use high contrast white noise stimuli to map neural receptive fields [25, 26].

Linear subunit RF responses to Gaussian noise are guaranteed to be Gaussian, so the lack of contrast normalization does not hurt performance in common laboratory conditions. With natural signals, the failure to normalize can hurt performance. Perhaps this is one reason why subunit models tend to generalize poorly to natural stimuli (but see [15]). It may be useful to incorporate response normalization in future instantiations of these models.

### Data-constrained Likelihood Functions

The class-conditional response distributions fully determine the likelihood function over the latent variable for any joint filter response **R** to an arbitrary stimulus. When the class-conditional response distributions are Gaussian, as they are here, the log-likelihood of latent variable value *X_u_* is quadratic in the encoding filter responses

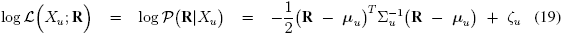

where 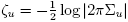. (Note that the likelihood function over the latent variable (Eq. 19) is distinct from likelihood function over the AMA-Gauss filters (Eq. 15).) Carrying out the matrix multiplication shows that the log-likelihood can be re-expressed as the weighted sum of squared, sum-squared (and linear) filter responses

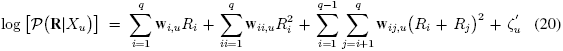

where *q* is the number of filters and where the weights are functions of the class-conditional mean and covariance for each value *X_u_* of the latent variable [6, 7]. specifically,

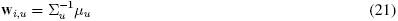

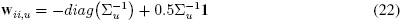

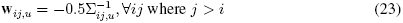

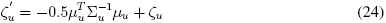

where *diag*(.) is a function that returns the diagonal of a matrix and **1** is a column vector of ones. (Note that in these equations *i* and *j* index different filters (see Fig. 2), not different latent variables and stimuli, as they do elsewhere in this manuscript). These equations (Eqs. 20 - 24) indicate that the log-likelihood of latent variable value *X_u_* is obtained by pooling the squared (and linear) responses of each receptive field with weights determined by the mean μ_*u*_ and covariance Σ_*u*_) of the subunit responses to stimuli with latent variable *X_u_*.

In the speed and disparity estimation tasks, nearly all of the information about the latent variable is carried by the class-conditional covariance; the covariance of the filter responses to natural stimuli changes significantly with changes in the latent variable (see Figs. 4.D-F, 5.EF, and 6.EF). Thus, the weights on the squared and the sum-squared filter responses change dramatically with the value of the latent variable (Fig. 9). In the speed estimation task, for example, the weights **w**_34_(*X*) on the sum-squared response of filter 3 and filter 4 peak at 0 deg/sec (see Fig 9.A). This peak results from the fact that the filter 3 and filter 4 response covariance is highest at 0 deg/sec (see Fig. 5.F; Eq.23). In contrast, very little information is carried by the class-conditional means; the mean filter responses to natural stimuli are always approximately zero. Hence, the weights on the linear subunit responses are approximately zero (see Eq. 21, Fig. 4.A-C).

**Figure 9:**
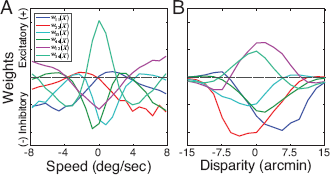
Quadratic pooling weights for computing the likelihood. Weights on squared and sum-squared filter responses (**w**_*ii*_(*X*) and **w**_*ij*_(*X*)) as a function of the latent variable. Weights on the linear filter responses are all approximately zero and are not shown. **A** Weights for speed estimation task. **B** Weights for disparity estimation task. Weights on squared responses are at maximum magnitude when the variance of the corresponding filter responses are at minimum. Weights on sum-squared responses are at maximum magnitude for latent variables yielding maximum response covariance (see Figs. 5.EF and 6.EF).

The filter response distributions determine the computations (i.e. quadratic pooling rules and weights) required to compute the likelihood of different latent variable values. If these computations (Eq. 20 - 24) are paired with an exponential output nonlinearity and implemented in a neuron, the neuron’s response 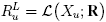 would represent the likelihood that a stimulus having a particular value *X_u_* of the latent variable elicited the observed filter responses **R**. This latent variable value *X_u_* would be the preferred stimulus of the likelihood neuron. We refer to this hypothetical neuron as an AMA-Gauss likelihood neuron (see Eq. 20).

Four example likelihood functions are shown in Fig. 10.A, one for each of four stimuli having a true speed of −4 deg/sec. Fig. 10.B shows four likelihood functions for stimuli having a true speed of 0 deg/sec. Fig. 10.C,D show likelihood functions for stimuli having −15 arcmin and 0 arcmin of binocular disparity, respectively. These plots show the likelihood functions, but they are not the standard way of assessing the response properties of neurons in cortex.

**Figure 10:**
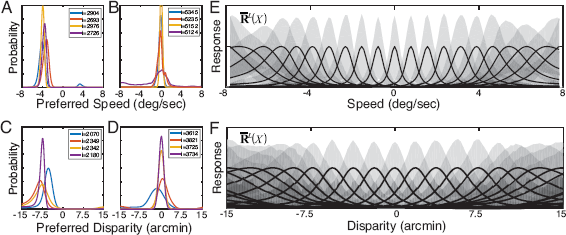
Likelihood functions for speed and disparity tasks. **A** Likelihood functions for four randomly chosen natural image movies having true speeds of 4 deg/sec. Each likelihood function represents the population response of the set of likelihood neurons, arranged by their preferred speeds. To ease the visual comparison, the likelihood functions are normalized to a constant volume by the sum of the likelihoods. **B** Same as A, but for movies with a true speed of 0 deg/sec. **C,D** Same as A,B but for stereo-images with −7.5 arcmin and 0.0 arcmin of disparity, respectively. **E** Tuning curves of speed-tuned likelihood neurons. For speeds sufficiently different from zero, tuning curves are approximately log-Gaussian and increase in width with speed. For near-zero speeds, tuning curves are approximately Gaussian. Each curve represents the mean response (i.e. tuning curve) of a likelihood neuron having a different preferred speed, normalized to a common maximum. Gray areas indicate 68% confidence intervals. **F** Tuning curves of disparity-tuned likelihood neurons.

The response properties of neurons in cortex are more commonly assessed by their tuning curves. Likelihood neuron tuning curves are obtained by first computing the mean likelihood neuron response across all natural stimuli having latent variable value *X_k_*

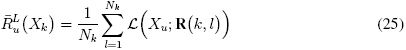

and then repeating for all values of the latent variable. Tuning curves for a population of likelihood neurons 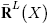 having a range of preferred speeds are shown in Fig. 10.E. The speed tuning curves are approximately Gaussian for preferred speeds near 0 deg/sec and approximately log-Gaussian otherwise. Consistent with these results, neurons in area MT have approximately log-Gaussian speed tuning curves, and have bandwidths that increase systematically with speed [33]. It is also interesting to note that while quadratic computations are required to optimally decode the latent variable directly from the filter responses (see Fig. 5.EF), likelihood neuron responses are linearly separable in speed. Similar points can be made about the disparity likelihood neurons (Fig. 10.F). The computations reported here thus constitute a general recipe for how to construct selective, invariant neurons having an arbitrary preferred stimulus (latent variable) from the responses of a small, well-chosen set of receptive fields.

### Linking AMA-Gauss and the Energy Model

Neural activity involved in many visual tasks has been productively modeled by energy-model-like (i.e. quadratic) computations [16, 13, 41]. We have shown that in two classic tasks (retinal speed and binocular disparity estimation), the class-conditional filter response distributions to natural stimuli are approximately Gaussian distributed. In such cases, quadratic combinations of the filter responses are the optimal computations and yield the likelihood of a particular value of the latent variable (Eq. 20). The weights are determined by the filter responses to natural stimuli (see Section 2). Thus, if these computations were instantiated in a neuron, then its response would represent the likelihood of latent variable (Fig. 2.C). The current results therefore constitute a normative explanation for why energy-model-like computations account for response properties of neurons involved in these tasks.

Interestingly, in recent years, discrepancies have emerged between the properties of neurons in cortex and the energy models that are often used to describe them [13, 44, 48]. Many of these discrepancies are a natural consequence of the optimal computations for estimating disparity and motion from natural stimuli[6, 7]. For example, the responses of motion- and disparity-selective neurons, have both been found to depend on multiple excitatory and suppressive subunit receptive fields, rather than the two exclusively excitatory subunit receptive fields posited by the energy model. Multiple subunit receptive fields have increased potential to select task-relevant information from each stimulus. Excitatory and inhibitory weighting schemes are required to use the selected information optimally. The quadratic computations in Eq. (20) specify exactly how to optimally weight and sum the responses from multiple receptive fields to achieve selectivity for particular latent variable values (also see Fig. 9). These computations yield more selective, invariant tuning curves (and improved estimation performance) than those of the standard energy model [6, 7], and follow directly from the normative framework employed here.

### Linking AMA-Gauss and the GQM: Connecting Normative and Response Triggered Analyses

In this section, we establish the formal similarities between AMA-Gauss and the Generalized Model (GQM), a recently developed subunit model for neural systems identification [40, 54]. The goal of the GQM is to identify (fit) the subunit receptive fields that drive a neuron’s response (Fig. 2.B). The goal of AMA-Gauss is to find the subunit receptive fields and quadratic pooling rules that are best for a particular task (Fig. 2.C). AMA can thus generate predictions about the subunit receptive fields that the GQM will recover from a neuron, under the hypothesis that the neuron computes the likelihood of a task relevant latent variable.

The GQM assumes that a neuron’s spiking or intra-cellular voltage response to a stimulus is given by

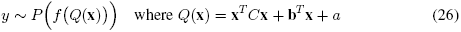

where *y* is the neural response, *P*(.)is the noise model, *f*(.) is a non-linearity, and λ = *f*(*Q*(**x**)) is the mean response. In [40], the authors use maximum likelihood methods to recover the parameters of the model given a set of stimuli, the neuron’s response to each stimulus, and a noise model. In AMA-Gauss, the log-likelihood of latent variable *X_u_* is given by

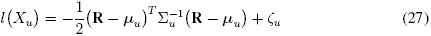

where μ_*u*_ and Σ_*u*_ are the class-conditional response mean and covariance and ζ_*u*_ is a constant. The noisy filter response vector **R** is given by the projection of the stimulus onto the filters **f** plus noise (Eqs. (4),(5)). Hence, Eq ( 27) can be rewritten as

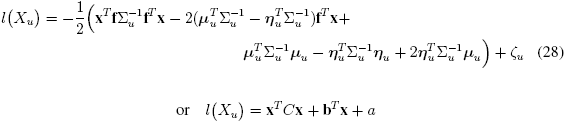

where 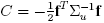 is a rank-*q* matrix where *q* is the number of filters, 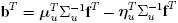, and 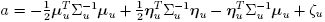. (Parameter values under the expected log-likelihood are provided in Appendix D). The parameters of the GQM are therefore simple functions of the AMA-Gauss encoding filters **f** and their responses to natural stimuli, conditional on latent variable *X_u_*. Given a hypothesis about the functional purpose of a neuron’s activity, AMA-Gauss could predict the parameters that the GQM would recover via response-triggered analyses.

The primary formal distinction between AMA-Gauss and the GQM is that AMA-Gauss explicitly models noise in the encoding filter responses, whereas the GQM models noise only after quadratic pooling of the filter responses; the GQM implicitly assumes noiseless filter responses. When subunit responses are noiseless, all subunit receptive fields spanning the same subspace (i.e. all linear filter combinations) provide an equivalent encoding. When responses are noisy (as they are in all biological systems), the stimulus encodings provided by different filters spanning the same subspace are no longer equivalent [8]. Future work will examine whether this distinction between AMA and the GQM can be leveraged to overcome a limitation common to all standard subunit models; namely, that their descriptions of neurons are unique only up to the subspace spanned by the subunit receptive fields (but see [27]).

## 4. Discussion

Accuracy Maximization Analysis (AMA) is a supervised Bayesian method for task-specific dimensionality reduction; it returns the encoding filters (receptive fields) that select the stimulus features that provide the most useful information about the task-relevant latent variable [21]. In conjunction with carefully collected databases of natural images and scenes and psychophysical experimental techniques, AMA has contributed to the development of ideal observers for several fundamental sensory-perceptual tasks in early- and mid-level vision [4, 6, 7, 21]. Unfortunately, AMA’s compute-time is high enough to render the method impractical for large problems without specialized computing resources.

We have developed AMA-Gauss, which makes the assumption that the class-conditional filter responses are Gaussian distributed and have shown that AMA-Gauss markedly reduces compute-time without compromising performance when the assumption is justified. We show that the assumption is justified for two fundamentally important visual tasks with natural stimuli (see Fig. 5 and Fig. 6, [6, 7]). These results provide a normative explanation for why energy model-like computations have proven useful in the study of motion and disparity estimation and discrimination. We speculate that the assumption will prove justified for other energy-model-related tasks in early vision (e.g. motion-in-depth estimation). AMA-Gauss also has the same formal structure as the Generalized Quadratic Model (GQM) a recently developed method for neural systems identification, raising the possibility that a single framework could be used both to predict and estimate the properties of involved in particular tasks.

There are several important implications of these results. First, the optimal filters and the optimal pooling rules for decoding the latent variable, are all determined by the properties of natural stimuli. If the training sets are representative of stimuli encountered in natural viewing, then the computations reported here should be optimal for the tasks of speed and disparity estimation. Second, at the right level of abstraction, the optimal solutions to these two different tasks share deep similarities, thereby raising the possibility that the same normative computational framework will apply to all energy-model related tasks.

### Response distributions: Gaussian vs. Heavy-tailed

The results reported here may appear to conflict with the widely reported finding that receptive field responses to natural images are highly non-Gaussian, with heavy tails sharp peaks at zero [18, 38, 10]). There are two explanations for this apparent discrepancy. First, previous analyses generally have not incorporated contrast normalization. Second, previous analyses are generally unsupervised and therefore do not condition on relevant latent variables (e.g. motion) [10]. Note that even when contrast normalization is incorporated and the class-conditional responses are Gaussian, the filter responses, marginalized over the latent variable, tend to be heavy-tailed because the marginals are mixtures of Gaussians 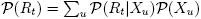 with different variances (see black curves in Fig. 5E,F and Fig. 6E,F). Therefore, our results are more similar to previous results than it may appear at first glance [43]. In general, heavy-tailed response distributions are yielded by response models that do not incorporate biologically plausible contrast normalization and response analyses that do not include latent variable conditionalization [51, 29]. Incorporating response normalization and latent variable conditionalization, as we have here, may help reveal statistical properties of receptive field responses to complex natural stimuli that has not yet been fully appreciated.

### Likelihood Functions:Data-Constrained vs Assumed

Evolution selects organisms because they perform certain critical sensory, perceptual, and behavioral tasks better than their evolutionary competitors. Certain features of sensory stimuli are more useful for some tasks than others. The stimulus features that are most useful to encode thus depend on the task-relevant latent variables that will be decoded from the stimuli. However, many models of neural encoding do not explicitly consider the tasks for which the encoded information will be decoded [38, 47] and many task-specific models of neural decoding do not explicitly consider how sensory stimuli and neural encoders constrain the information available for decoding [17, 30].

The approach advanced here is an early attempt to address both issues simultaneously. By performing task-specific analyses using thousands of individual natural stimuli, learning the optimal filters, and characterizing the class-conditional responses to natural stimuli, we determined the likelihood functions that optimize performance in natural viewing. The likelihood functions that result from the filter response distributions are (on average) log-Gaussian in speed and disparity, with widths that increase with the value of the latent variable. In previous work with natural stimuli, we showed that the optimal receptive fields, response distributions, and resulting likelihood functions are robust to significant variation in the shape of the prior, cost function, and noise power [8]. It is reasonable to conclude that the task and the constraints imposed by natural stimuli are the most important determinants of the width and shape of the likelihood functions.

Some prominent theories of neural processing operate on the assumption that likelihood functions can take on arbitrary widths and shapes via flexible allocation of neural resources [19, 22, 46, 52]. Some reports have gone further to suggest that, in the context of Bayesian efficient coding, the prior probability distribution over the latent variable is the primary factor determining the widths and shapes of the likelihood functions [19, 52]. These reports predict that if the prior probability distribution is flat, the likelihood functions will be symmetric and have widths that remain constant with changes in the value of the latent variable. These reports also predict that if the prior probability distribution is non-uniform (e.g. peaked at zero), the likelihood functions will be asymmetric with widths that change systematically with the latent variable.

In the tasks that we examined, we found that asymmetric likelihood functions optimize performance despite the fact that the training sets from which the optimal filters were learned had flat priors over the latent variable (see Results; [6, 7, 8]). These results appear at odds with the predictions of previous reports. However, these previous reports do not model the impact of natural stimulus variation. The implicit assumption is that task-irrelevant (’nuissance’) stimulus variation can be ignored [19, 52]. If the goal is to understand optimal task-specific processing of natural signals, our results indicate that such variation cannot be ignored, Indeed, task-relevant and irreducible task-irrelevant natural stimulus variability are almost certainly the most important determinants of likelihood function shapes and widths.

In natural viewing, visual estimates are driven primarily by stimulus measurements (likelihood functions), not by prior distributions. If estimates were driven only by the prior, observers could not respond to spatial or temporal changes in the environment. A full account of task-specific perceptual processing and its underlying neurophysiology must therefore incorporate natural stimulus variability. Future studies on the efficient allocation of neural resources should verify that the likelihood functions used in modeling efforts can be constructed by nervous systems given the constraints imposed by natural stimuli.

### Natural vs. Artificial Stimuli

The problem of estimating speed and disparity from natural images is different from the problem of estimating speed and disparity with artificial laboratory stimuli in at least one important respect. Variability amongst natural stimuli having the same latent variable level is typically greater than variability amongst artificial stimuli commonly used in vision and visual neuroscience experiments. In motion experiments (Fig. 11.A), drifting Gabors and random-dot kinematograms are common artificial stimuli. In disparity experiments, phase-shifted binocular Gabors and random-dot stereograms are common artificial stimuli (Fig. 11.B). The statistical properties of these artificial stimuli are notably different than the statistical properties of natural stimuli. Gabors have Gaussian amplitude spectra and random-dot stereograms have delta auto-correlation functions. Natural stimuli have a rich variety of textures and shapes that cause significant variation in their 1/f amplitude spectra and auto-correlation functions.

**Figure 11:**
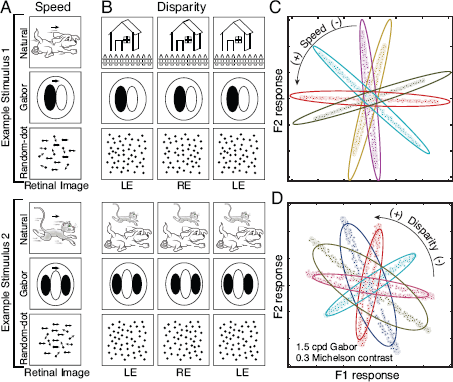
Natural stimuli, artificial stimuli, and class-conditional responses. Many different retinal images are consistent with a given value of the task-relevant latent variable. These differences cause within-class (task-irrelevant) stimulus variation. Within-class stimulus variation is greater for natural stimuli than for typical artificial stimuli used in laboratory experiments. **A** Stimuli for speed estimation experiments. Two different example stimuli are shown for each stimulus type: natural stimuli (represented by a cartoon line-drawings), Gabor stimuli, and random-dot stimuli. Both example stimuli for each stimulus type drift at exactly the same speed, but create different retinal images. Natural stimuli cause more within-class retinal stimulus variation than artificial stimuli. **B** Same as A, but for disparity. **C** Speed task: class-conditional responses to contrast-fixed 1.0 cpd drifting Gabors with random phase (speed task). Colors indicate different speeds. Ellipses represent filter responses to natural stimuli having the same speeds. **D** Disparity task: Class-conditional responses to contrast-fixed 1.5 cpd binocular Gabors with random phase. Class-conditional responses no longer have Gaussian structure, and instead have ring structure.

To examine the impact of artificial stimuli on the class-conditional responses, we created artificial stimulus sets comprised of contrast-fixed, phase-randomized Gabors drifting at different speeds and having different amounts of disparity. For each task, the spatial frequency of the carrier was closely matched to the preferred spatial frequency of the first two optimal filters (1.0 cpd for speed, 1.5 cpd for disparity). Joint filter responses to these artificial stimuli are shown in Fig. 11.C,D; they are notably different than the filter responses to natural stimuli. Although the class-conditional responses to Gabors are approximately aligned with the major axis of the Gaussian characterizing responses to corresponding natural stimuli, the responses themselves are no longer Gaussian distributed, exhibiting ring-shaped structure instead. Thus, determining the optimal rules for processing natural stimuli by analyzing only artificial stimuli is likely to be a difficult enterprise.

These results suggest another conclusion that may be somewhat counterintuitive given the history of the field. The tradition in vision science has been to eliminate irrelevant stimulus variation from experimental protocols by using simple artificial stimuli. These stimuli are easy to characterize mathematically and manipulate parametrically. But artificial stimuli lack the richness and variability that visual systems evolved to process. Analyzing complex, variable natural stimuli may reveal simple (e.g. Gaussian) statistical structure that might otherwise be missed. We believe that the results presented here highlight the importance of conducting rigorous, well-controlled, task-focused computational and behavioral investigations with natural stimuli. These investigations complement classic studies with artificial stimuli, and provide a fuller picture of how visual systems function in natural circumstances.

### Limitations and Future Directions

The results presented here represent the first in what we hope is a series of steps to link normative models for natural tasks and descriptive models of neural response. However, while we believe that developing AMA-Gauss and demonstrating its links to methods for neural systems identification are useful advances, several limitations should be kept in mind. Here, we address the drawbacks of the natural stimulus sets, the general applicability of AMA-Gauss, and the importance of the links that we have drawn to descriptive models of neural response.

The natural image sets used in this manuscript had natural contrast distributions and photographic textures, but they lacked natural depth structure. All motion signals were rigid and all disparity signals were planar. Future work will examine the impact of non-rigid motion (e.g. looming) and local depth variation (e.g. occlusion) on performance. We have recently collected a dataset of stereo-images that addresses this limitation [9]. Each stereo-image has co-registered distance data from which groundtruth disparity patterns can be computed. Pilot analyses suggest that the results presented in the current manuscript will hold for natural stereo-images with local depth variation. We suspect, but we are not yet well-positioned to show, that the same will be true of motion signals having natural depth variation.

AMA-Gauss is the appropriate normative framework for understanding energy-model-related tasks, but the general usefulness of AMA-Gauss is unknown. AMA-Gauss makes the best possible use of the first- and second-order filter response statistics, but it is blind to higher-order response statistics that exist in natural motion [32] and natural disparity signals. To increase generality, one could develop a variant of the method that incorporates rectification into the response model. This modification would confer the ability, at least in principle, to pick up on potentially useful higher-order motion and disparity cues, and provide a normative model that complements other methods for neural systems identification [31].

## 5. Conclusion

In this paper, we develop AMA-Gauss, a new form of AMA that incorporates the assumption that the class-conditional filter responses are Gaussian distributed. We use AMA-Gauss to establish links between task-specific normative models of speed and disparity estimation and the motion- and disparity-energy models, two popular descriptive models of neurons that are selective for those quantities. Our results suggest that energy-model-like (i.e. quadratic) computations are optimal for these tasks in natural scenes. We also establish the formal similarities between AMA-Gauss and the Generalized Quadratic Model (GQM), a recently developed model for neural systems identification. The developments presented here forge links between normative task-specific modeling and powerful statistical tools for describing neural response, and demonstrate the importance of analyzing natural signals in perception and neuroscience research.

## Appendix A Gradient of the Likelihood Function

In any given training set having *N* stimuli, each stimulus is associated with some category *k* and an associated stimulus from that category *l*. Let us denote this pair (*k*, *l*)for the *i^th^* sample point with (*k_i_*, *l_i_*). Then assuming that the response distribution conditioned on the classes is Gaussian, the likelihood function can be written as

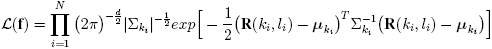

Substituting the expression for the noisy responses (Eq. (5)) and defiing 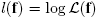 yields the log-likelihood function of the AMA-Gauss filters

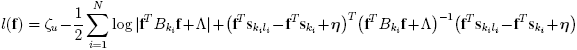

where 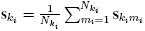 and 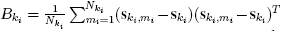 are the class-conditional stimulus mean and covariance matrix, respectively, and 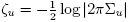 is a constant.

Rearranging to segregate terms that do not depend on noise samples

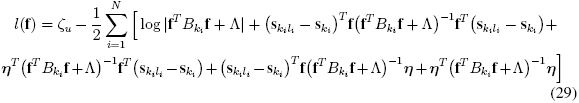

where **f**^*T*^ *B***f** + Λ is a symmetric matrix. Recognizing that each term in Eq 29 is a scalar, and rewriting using the properties that *Tr*(*a*)= *a*, *Tr*(**AB**) = *Tr*(**BA**) and *Tr*(**A**) = *Tr*(**A**^*T*^) yields

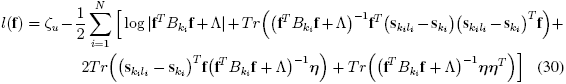

To determine the gradient of the log-likelihood 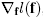, we derive the gradient of each term in Eq. 30 separately below. Before doing so, we state some standard matrix results that will be used in the derivation [42].

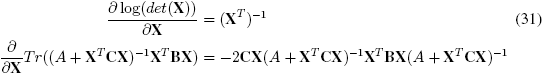

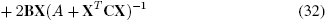

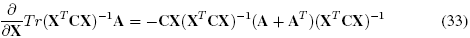

The gradient of the first term in Eq 30 is obtained by using Eq 31 and the chain rule of differentiation

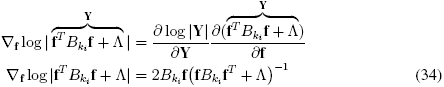

The gradient of the second term in Eq 30 is obtained using Eq 32

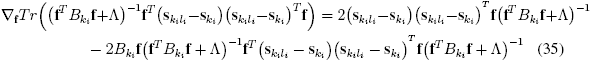

The gradient of the third term is obtained using Eq 33 and the chain rule of differentiation

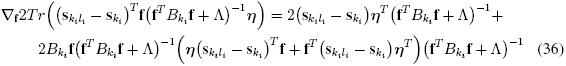

The gradient of the fourth term is similarly obtained using Eq 33

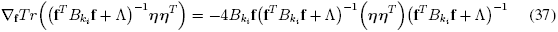

The full gradient of the AMA-Gauss filter log-likelihood *l*(**f**) stated in Eq 30 can therefore be found by combining Eqs 34-37.

The gradient of the expected log-likelihood follows directly from the gradient of the log-likelihood. The response noise 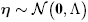 is normally distributed (Eq 6); therefore, 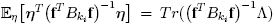. Substituting into Eq 30 yields the expected log-likelihood of the AMA-Gauss filters

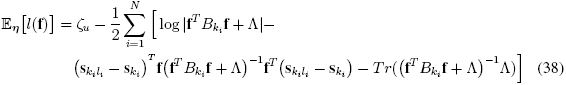

The gradient of the expected log-likelihood, using Eqs 34,35, and 37, is given by

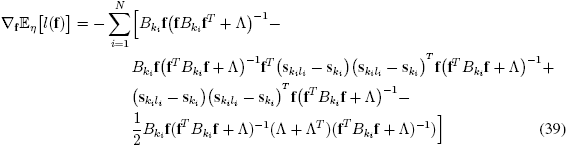

## Appendix B Gradient of *L*_2_ cost function

The average expected cost across all the stimuli is

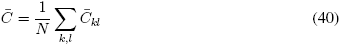

Given the squared error loss function, the expected cost per stimuli can be written as

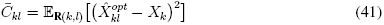

where 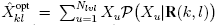 since the optimal estimate for a squared error function is the mean of the posterior, i.e. 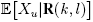. Using the approximation that the expected cost of each stimulus is equal to the cost given the expected response [21] yields

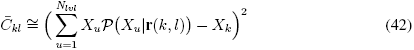

Therefore, to evaluate the gradient of the total cost we just need to evaluate the expression for the gradient of the expected cost of each stimulus. Hence,

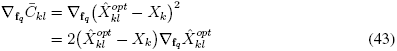

The gradient of the optimal estimate given the mean response is

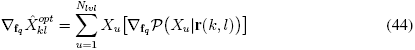

Hence, the problem reduces to finding 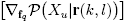

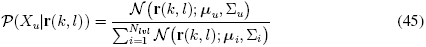

Making substitutions in Eq (45) gives

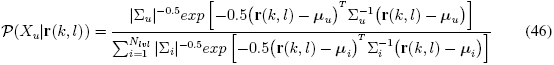

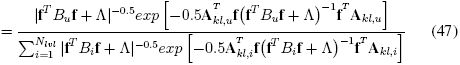

where **A**_*kl,u*_=**s**_*kl*_ − **s**_*u*_. The gradient of the posterior probability can then be evaluated using the following relation with the gradient of the logarithm of the posterior probability

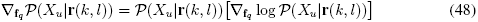

Taking the natural logarithm of the posterior yields

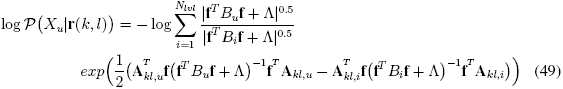

Next, we define new variables to simplify this expression for the log posterior probability and the subsequent derivation of its gradient. Let each term in the summation in Eq (49) be

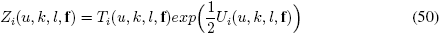

where 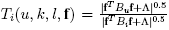 is the scale factor in each term in the summation in Eq (50) and where 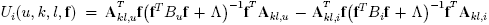 is the exponentiated term in each term of the sum in Eq (50). Hence, by substituting Eq. (50) into Eq (49) the simplified expression for the log posterior is

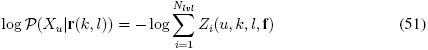

The gradient of the log posterior probability can therefore be expressed as

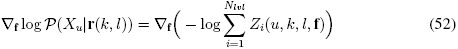

The gradient of the log is

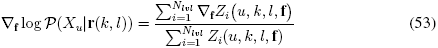

Expanding the numerator by substituting Eq (50) using the chain rule for differentiation

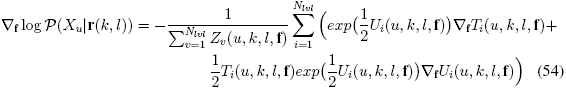

The remaining terms to be evaluated are 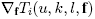 and 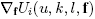.

The expression for 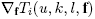 is

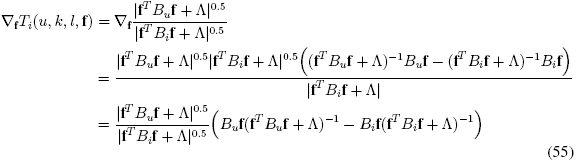

The expression for 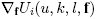 is

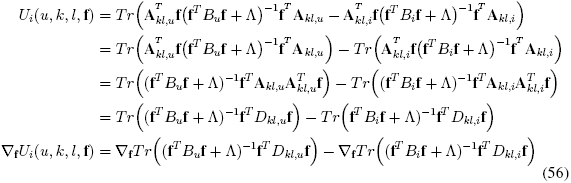

where 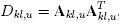. The expression for the gradient of the trace in Eq (56) is obtained by using Eqs (32). Thus,

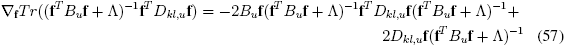

The gradient 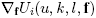 is obtained by substituting Eq (57) into Eq (56). The gradient of 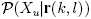 is obtained by substituting Eq (55) and Eq (56) into Eq (54). The gradient of the posterior probability is obtained by plugging Eq (54) into Eq (48). The gradient of the cost for each stimulus is obtained by plugging Eq (48) into Eq (44), and then plugging that result into Eq (43)

## Appendix C AMA-Gauss Gradient with *L*_0_ / KL-divergence cost function

The total cost for a set of filters is given by the average expected cost across all stimuli

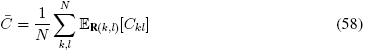

Given the 0,1 cost function, the cost associated with the filter response to an arbitrary stimulus is given by 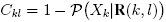. This cost is monotonic with KL-divergence and we refer to this cost as the KL-cost.

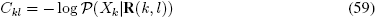

We approximate the expected cost associated with each stimulus with the expected cost given the mean response [21]. Thus, we have

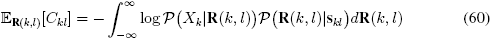

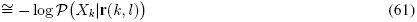

Therefore, the total cost for a set of filters is given by

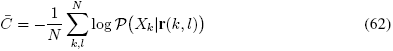

Hence, the gradient of the total expected cost 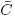 can then be written as

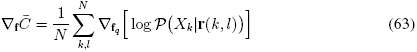

The full expression for the expected cost 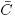 is obtained by substituting the expression for 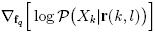 given by Eq (54), Eq (55), and Eq (56) in Appendix (B).

## Appendix D Connection between AMA-Gauss and GQM

The log-likelihood of latent variable *X_u_* using Eq (28) can be written as

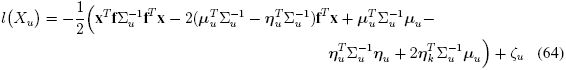

where 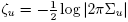 is a constant. The expected log-likelihood can then be written as

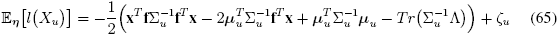

It is evident from Eq (65) that 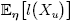 is of the form 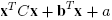 where

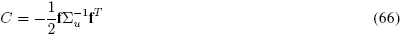

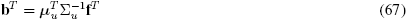

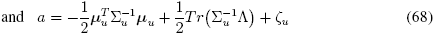

## Footnotes

1. AMA-Gauss software (Matlab) is available at https://www.github.com/burgelab/AMA

## References

[1] E H Adelson and J R Bergen. Spatiotemporal energy models for the perception of motion. JOSA A, 2(2):284–299, 1985.

[2] D G Albrecht and W S Geisler. Motion selectivity and the contrast-response function of simple cells in the visual cortex. Visual neuroscience, 7(06):531–546, 1991.

[3] A J Bell and T J Sejnowski. The “independent components” of natural scenes are edge filters. Vision research, 37(23):3327–3338, 1997.

[4] J Burge and W S Geisler. Optimal defocus estimation in individual natural images. Proceedings of the National Academy of Sciences, 108(40):16849–16854, 2011.

[5] J Burge and W S Geisler. Optimal defocus estimates from individual images for autofocusing a digital camera. In Proceedings of the IS&T/SPIE 47th Annual Meeting. Proceedings of SPIE, 2012.

[6] J Burge and W S Geisler. Optimal disparity estimation in natural stereo images. Journal of vision, 14(2):1–1, 2014.

[7] J Burge and W S Geisler. Optimal speed estimation in natural image movies predicts human performance. Nature communications, 6, 2015.

[8] J Burge and P Jaini. Accuracy Maximization Analysis for Sensory-Perceptual Tasks: Computational Improvements, Filter Robustness, and Coding Advantages for Scaled Additive Noise. PLoS computational biology, 13(2):e1005281, 2017.

[9] J Burge, B C McCann, and W S Geisler. Estimating 3d tilt from local image cues in natural scenes. Journal of Vision, 16(13):2–2, 2016.

[10] C F Cadieu and B A Olshausen. Learning intermediate-level representations of form and motion from natural movies. Neural computation, 24(4):827–866, 2012.

[11] R D Cook and L Forzani. Likelihood-based sufficient dimension reduction. Journal of the American Statistical Association, 104 (485):197–208, 2009.

[12] RD Cook, L Forzani, and AF Yao. Necessary and sufficient conditions for consistency of a method for smoothed functional inverse regression. Statistica Sinica, pages 235–238, 2010.

[13] BG Cumming and GC DeAngelis. The physiology of stereopsis. Annual review of neuroscience, 24(1):203–238, 2001.

[14] G C DeAngelis. Seeing in three dimensions: the neurophysiology of stereopsis. Trends in cognitive sciences, 4(3):80–90, 2000.

[15] M Eickenberg, R J Rowekamp, M Kouh, and T O Sharpee. Characterizing responses of translation-invariant neurons to natural stimuli: maximally informative invariant dimensions. Neural computation, 24(9):2384–2421, 2012.

[16] R C Emerson, J R Bergen, and E H Adelson. Directionally selective complex cells and the computation of motion energy in cat visual cortex. Vision research, 32(2):203–218, 1992.

[17] M O Ernst and M S Banks. Humans integrate visual and haptic information in a statistically optimal fashion. Nature, 415(6870):429-433, 2002.

[18] D J Field. Relations between the statistics of natural images and the response properties of cortical cells. Journal of the Optical Society of America. A, Optics and image science, 4(12):2379–2394, 1987.

[19] D Ganguli and E P Simoncelli. Efficient sensory encoding and bayesian inference with heterogeneous neural populations. Neural computation, 2014.

[20] W S Geisler and D G Albrecht. Visual cortex neurons in monkeys and cats: detection, discrimination, and identification. Visual neuroscience, 14(5):897–919, 1997.

[21] W S Geisler, J Najemnik, and A D Ing. Optimal stimulus encoders for natural tasks. Journal of vision, 9(13):17–17, 2009.

[22] A R Girshick, M S Landy, and E P Simoncelli. Cardinal rules: visual orientation perception reflects knowledge of environmental statistics. Nature neuroscience, 14(7):926–932, 2011.

[23] D J Heeger. Normalization of cell responses in cat striate cortex. Visual neuroscience, 9(02):181–197, 1992.

[24] H Hotelling. Analysis of a complex of statistical variables into principal components. Journal of educational psychology, 24(6):417, 1933.

[25] J P Jones and L A Palmer. An evaluation of the two-dimensional gabor filter model of simple receptive fields in cat striate cortex. Journal of neurophysiology, 58(6):1233–1258, 1987.

[26] J P Jones and L A Palmer. The two-dimensional spatial structure of simple receptive fields in cat striate cortex. Journal of neurophysiology, 58(6):1187–1211, 1987.

[27] J Kaardal, J D Fitzgerald, M J Berry, and T O Sharpee. Identifying functional bases for multidimensional neural computations. Neural computation, 2013.

[28] M S Lewicki. Efficient coding of natural sounds. Nature neuroscience, 5(4):356–363, 2002.

[29] S Lyu and E P Simoncelli. Modeling multiscale subbands of photographic images with fields of Gaussian scale mixtures. Pattern Analysis and Machine Intelligence, IEEE Transactions on, 31(4):693–706, 2009.

[30] W J Ma, Jeffrey M Beck, Peter E Latham, and A Pouget. Bayesian inference with probabilistic population codes. Nature neuroscience, 9(11):1432–1438, 2006.

[31] J M McFarland, Y Cui, and D A Butts. Inferring nonlinear neuronal computation based on physiologically plausible inputs. PLoS computational biology, 9(7):e1003143, 2013.

[32] E I Nitzany and J D Victor. The statistics of local motion signals in naturalistic movies. Journal of vision, 14(4), 2014.

[33] H Nover, C H Anderson, and G C DeAngelis. A logarithmic, scale-invariant representation of speed in macaque middle temporal area accounts for speed discrimination performance. The Journal of neuroscience : the official journal of the Society for Neuroscience, 25(43):10049–10060, 2005.

[34] I Ohzawa. Mechanisms of stereoscopic vision: the disparity energy model. Current opinion in neurobiology, 8(4):509–515, 1998.

[35] I Ohzawa, G C DeAngelis, and R D Freeman. Stereoscopic depth discrimination in the visual cortex: neurons ideally suited as disparity detectors. Science, 249(4972):1037–1041, 1990.

[36] I Ohzawa, G C Deangelis, and R D Freeman. Encoding of binocular disparity by complex cells in the cat’s visual cortex. Journal of neurophysiology, 77(6):2879–2909, 1997.

[37] B A Olshausen and Field. Emergence of simple-cell receptive field properties by learning a sparse code for natural images. Nature, 381 (6583):607–609, 1996.

[38] B A Olshausen and D J Field. Sparse coding with an overcomplete basis set: A strategy employed by v1? Vision research, 37(23):3311–3325, 1997.

[39] M Pagan, E P Simoncelli, and N C Rust. Neural Quadratic Discriminant Analysis: Nonlinear Decoding with V1-Like Computation. Neural computation, pages 1–29, 2016.

[40] I M Park, E W Archer, N Priebe, and J W Pillow. Spectral methods for neural characterization using generalized quadratic models. In Advances in neural information processing systems, pages 2454–2462, 2013.

[41] Q Peng and B E Shi. The changing disparity energy model. Vision research, 50(2):181–192, 2010.

[42] K B Petersen, M S Pedersen, et al. The matrix cookbook. Technical University of Denmark, 7:15, 2008.

[43] D L Ruderman and W Bialek. Statistics of natural images: Scaling in the woods. Physical review letters, 73(6):814, 1994.

[44] N C Rust, O Schwartz, J A Movshon, and E P Simoncelli. Spatiotemporal elements of macaque v1 receptive fields. Neuron, 46 (6):945–956, 2005.

[45] O Schwartz, J W Pillow, N C Rust, and E P Simoncelli. Spike-triggered neural characterization. Journal of vision, 6(4):484–507, 2006.

[46] H S Seung and H Sompolinsky. Simple models for reading neuronal population codes. Proceedings of the National Academy of Sciences, 90(22):10749–10753, 1993.

[47] E P Simoncelli and B A Olshausen. Natural image statistics and neural representation. Annual review of neuroscience, 24(1):1193-1216, 2001.

[48] S Tanabe, R M Haefner, and B G Cumming. Suppressive mechanisms in monkey v1 help to solve the stereo correspondence problem. Journal of Neuroscience, 31(22):8295–8305, 2011.

[49] M E Tipping and C M Bishop. Probabilistic principal component analysis. Journal of the Royal Statistical Society: Series B (Statistical Methodology), 61(3):611–622, 1999.

[50] B Vintch, J A Movshon, and E P Simoncelli. A convolutional subunit model for neuronal responses in macaque v1. The Journal of Neuroscience, 35(44):14829–14841, 2015.

[51] Z Wang, A C Bovik, H R Sheikh, and E P Simoncelli. Image quality assessment: from error visibility to structural similarity. Image Processing, IEEE Transactions on, 13(4):600–612, 2004.

[52] X Wei and A Stocker. A bayesian observer model constrained by efficient coding can explain’anti-bayesian’percepts. Nature neuroscience, 18(10):1509–1517, 2015.

[53] D R Williams. Visibility of interference fringes near the resolution limit. JOSA A, 2(7):1087–1093, 1985.

[54] A Wu, I M Park, and J W Pillow. Convolutional spike-triggered covariance analysis for neural subunit models. In Advances in Neural Information Processing Systems, pages 793–801, 2015.

